## Supplementary Information for "Emergence of ion-channel mediated electrical oscillations in *Escherichia coli* biofilms"

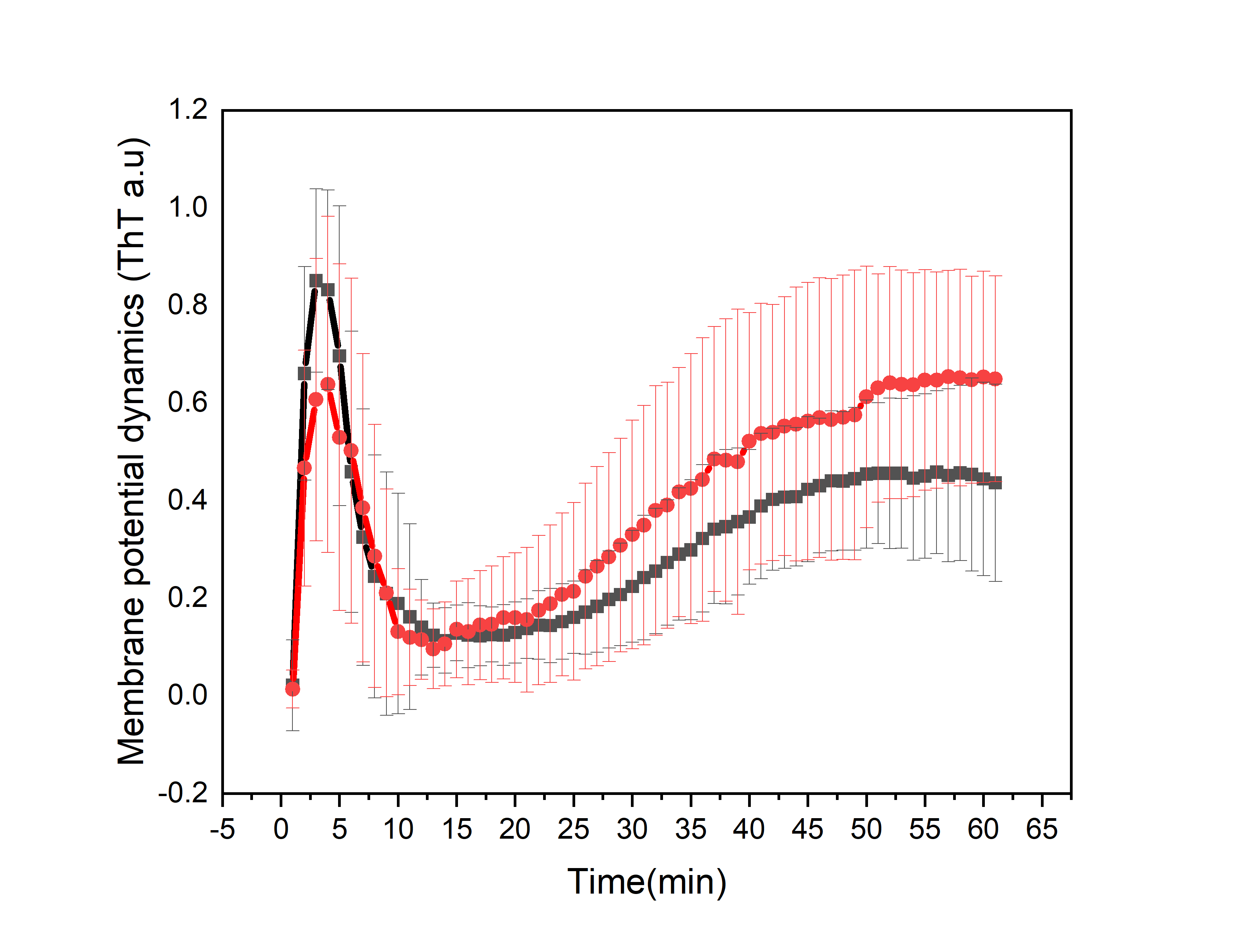

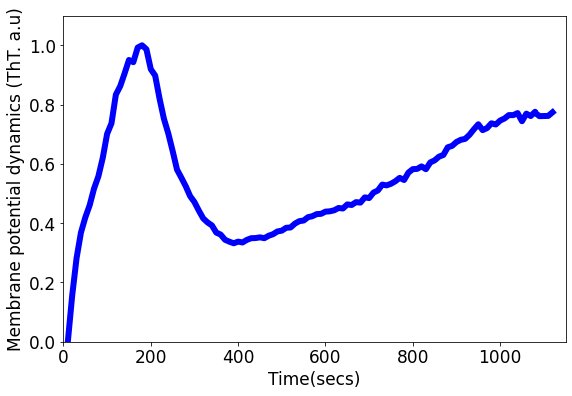

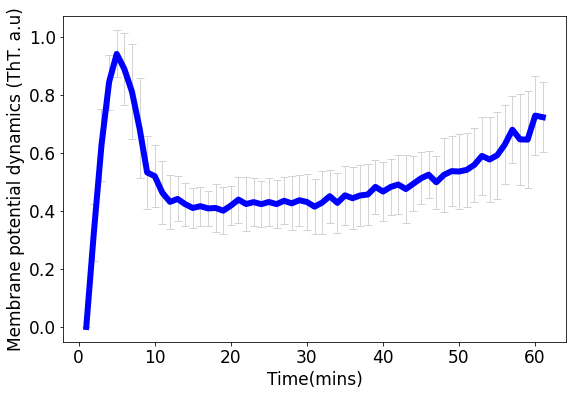

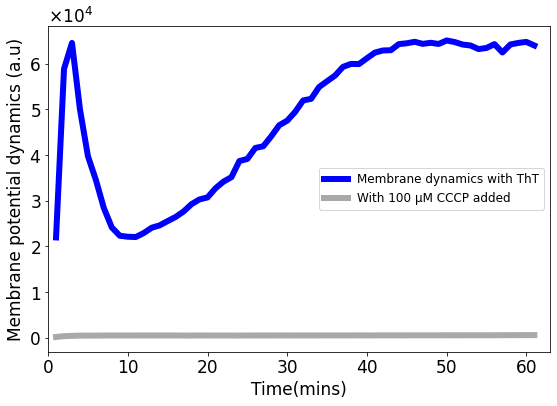

**A**

**B**

**C**

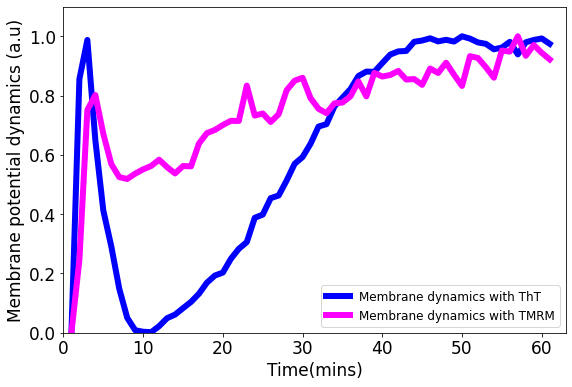

**D**

**E**

**Supplementary Fig 1**. (A) ThT fluorescence as a function of time shows the *E. coli* membrane potential dynamics under light stress with images collected every 10 seconds. (B) ThT fluorescence as a function of time shows the membrane potential dynamics for both wildtype *E. coli* strains, DH5α (red) and *E. coli* BW25113 (black) (Data: mean±SD). (C) ThT fluorescence as a function of time shows the membrane potential dynamics of *E. coli* in M9 media. The light stress was applied continuously each minute for 1 hour with images collected every minute. (D) TMRM and ThT fluorescence as a function of time with *E. coli*. The cationic dye, TMRM is used to confirm the membrane potential dynamics with ThT. (E) ThT fluorescence as a function of time. CCCP quenches membrane potential dynamics. CCCP was added to cell suspension and left for 50 minutes before exposure to blue light stress.

$$0.54 mm$$

Media inlet

To reservoir

**60x**

$$\boldsymbol{Blue light}$$

$$\boldsymbol{Objective}$$

$$\boldsymbol{Mature biofilm}$$

**A**

**B**

$$\boldsymbol{To reservoir}$$

$$\boldsymbol{To media pump}$$

$$\boldsymbol{Six flow channels}$$

$$\boldsymbol{tubes}$$

$$\boldsymbol{connector}$$

$$\boldsymbol{direction of flow}$$

$$25.5 mm$$

$$75.5 mm$$

**Supplementary Fig 2**. Schematic of the IBIDI flow cells used for all the experiments. (A) IBIDI uncoated glass bottom µ-Slide VI^0.5^ showing the inlet and outlet openings. The 0.54 mm depth enabled growth of biofilms in 3D. (B) IBIDI uncoated glass bottom µ-Slide VI^0.5^ with six identical wells for the growth of *E. coli* biofilms.

**A**

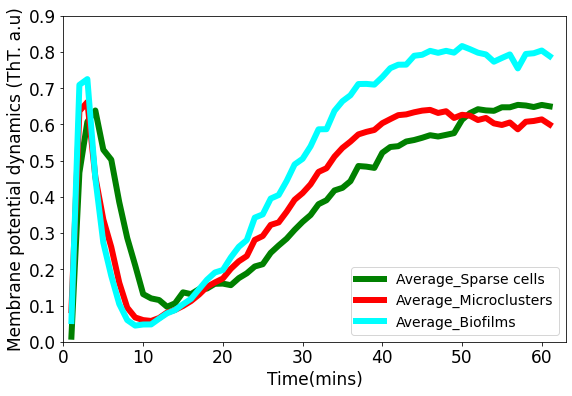

**B**

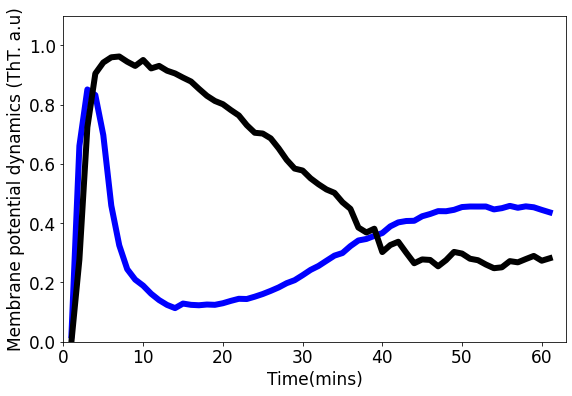

**C**

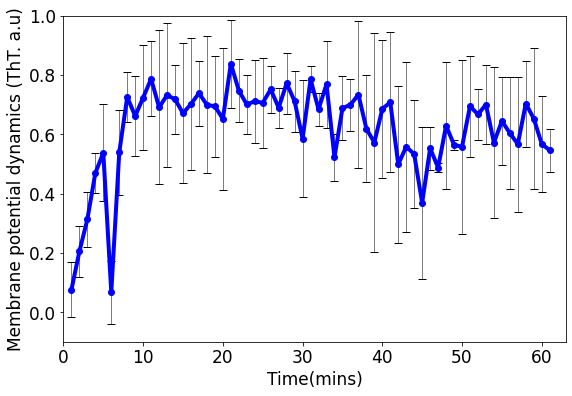

**D**

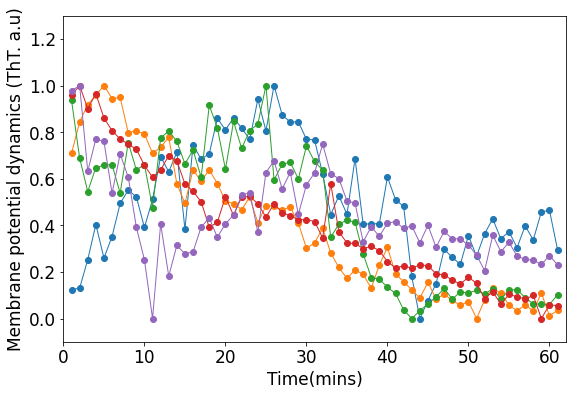

**E**

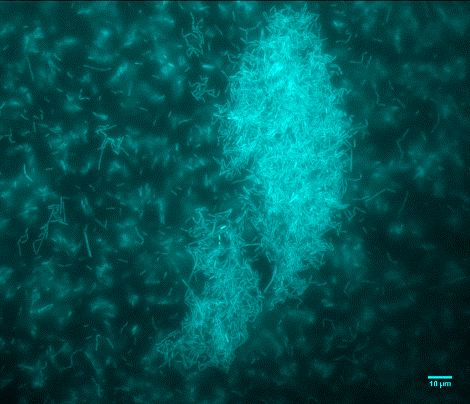

**Supplementary Fig 3:** A) Representative fluorescence image of a mature sessile biofilm labelled with ThT. The scale bar is 10 µm. (B) ThT fluorescence as a function of time for sparse cells, clustered cells and biofilms. First peak latency is much shorter in *E. coli* biofilms than in single cells and microclusters. (C) ThT fluorescence shown as a function of time of irradiation. Deletion of Kch inactivates the second peak in *single cell* *E. coli* BW25113 ∆*kch*-mutants. Data is a mean from three experimental replicates per time point for *E. coli* BW25113 ∆*kch*-mutants (black) and BW25113 (blue). (D) Membrane potential dynamics of kch-complemented DH5α. (E) Calcium flux in single cells DH5α under light stress.

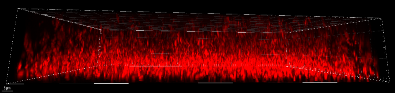

**1 min**

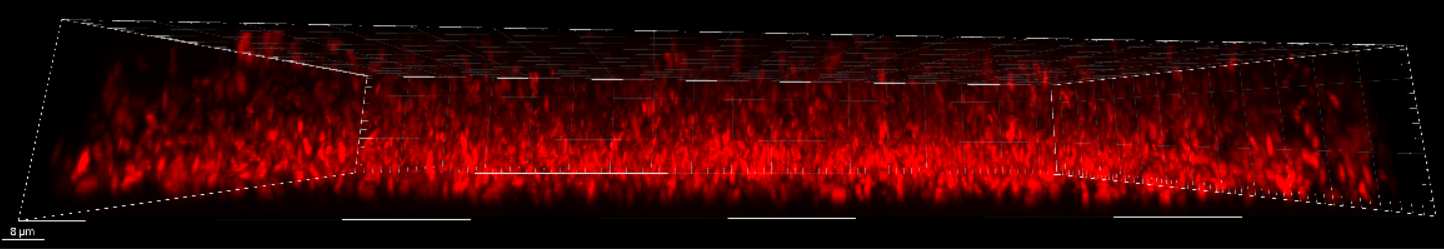

**3 min**

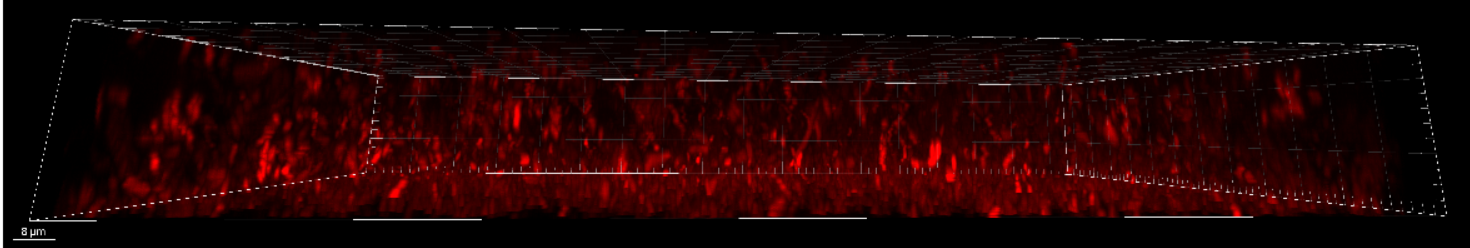

**10 min**

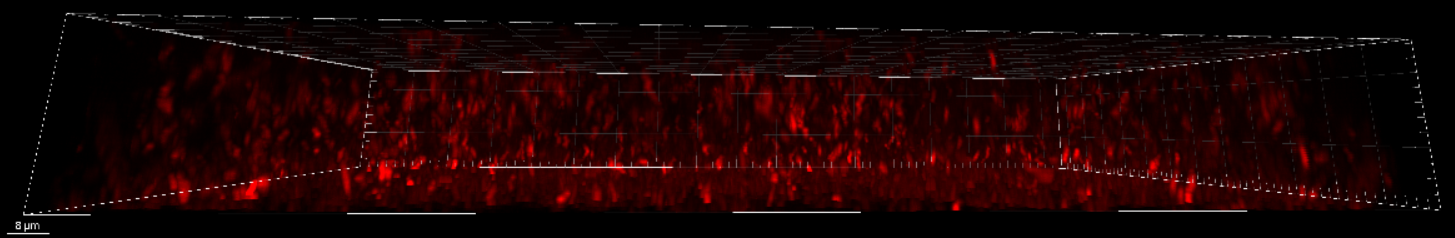

**15 min**

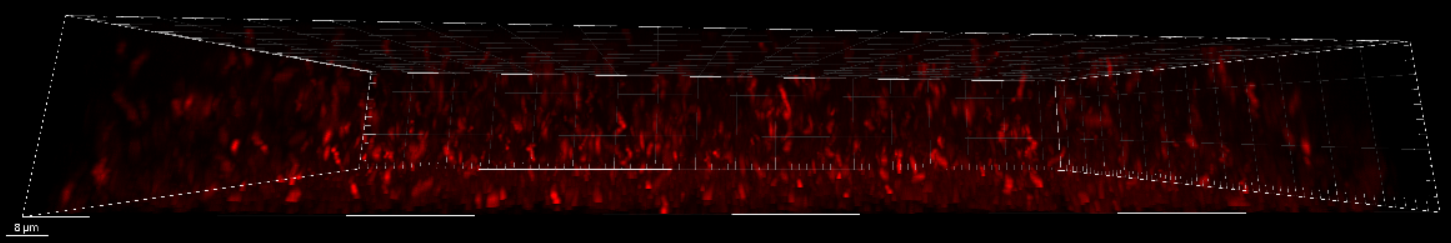

**20 min**

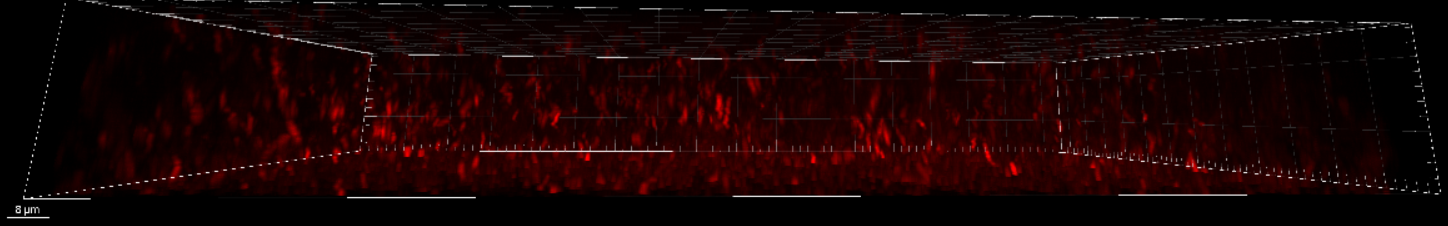

**30 min**

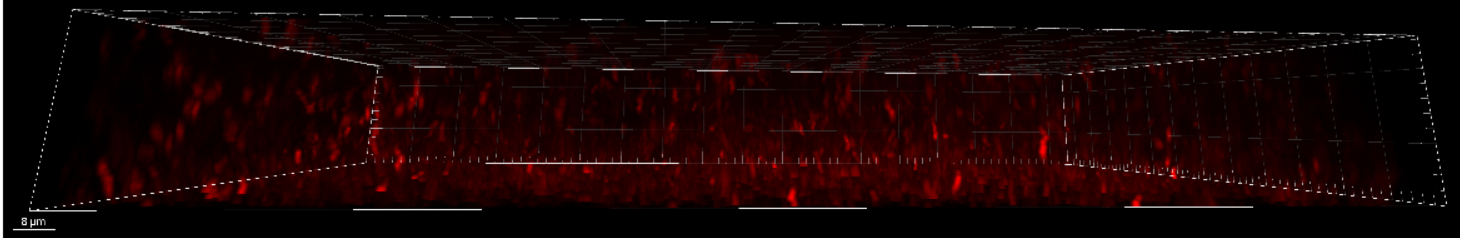

**40 min**

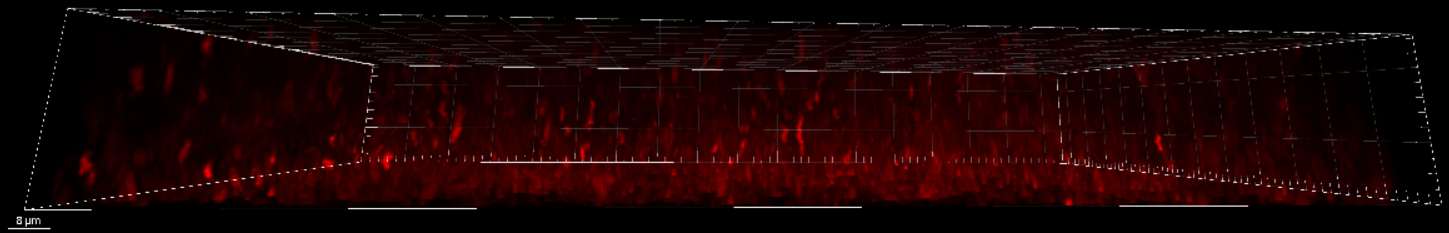

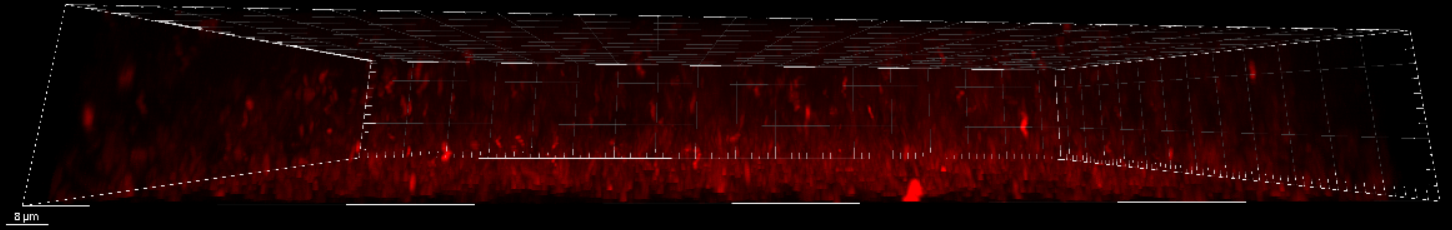

**50 min**

**60 min**

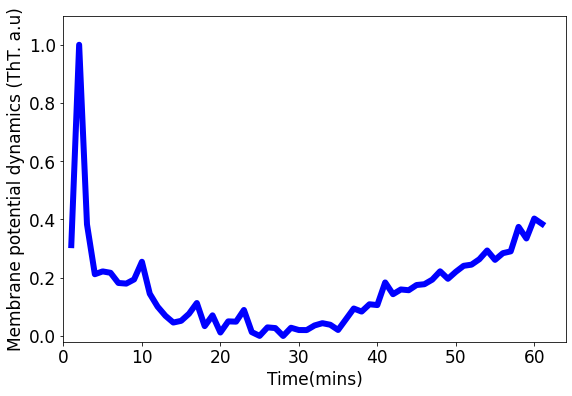

**B**

**A**

**Supplementary Fig 4:** (A) Timelapse of the sagittal section of the three-dimensional *E. coli* biofilm obtained using the confocal microscope. (B) Representative global ThT intensity membrane potential dynamics trace as a function of time obtained from a 3D DH5α *E. coli* biofilm. The scale bars for all the image are 8 µm.

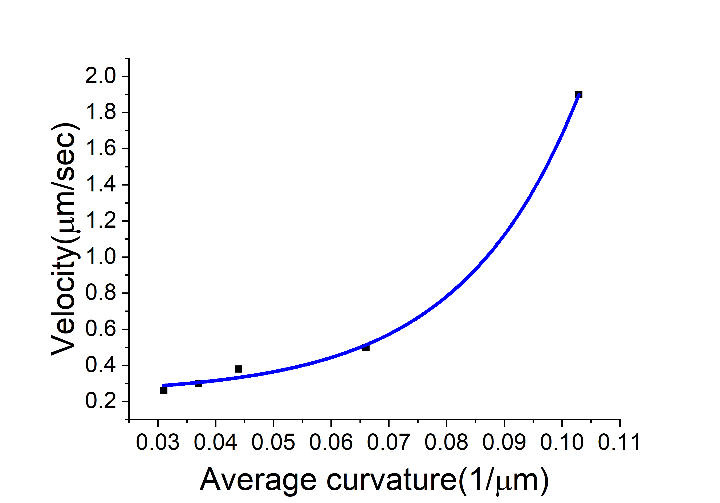

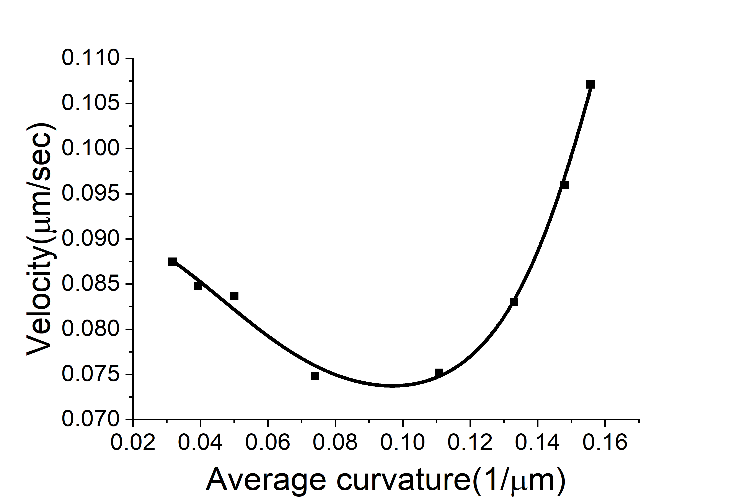

**II**

**I**

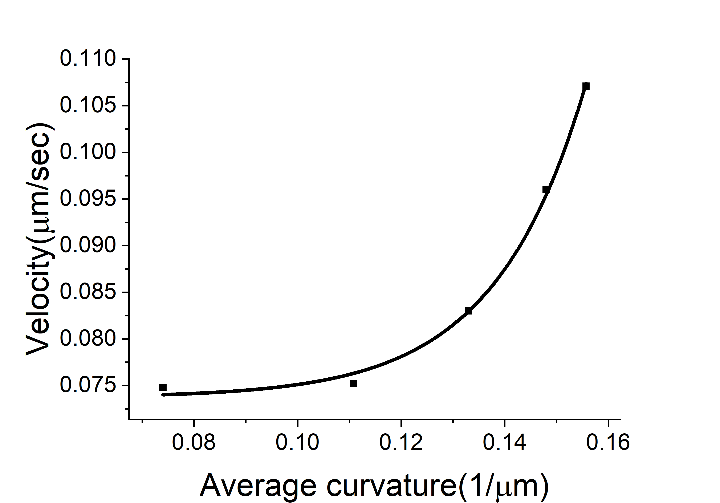

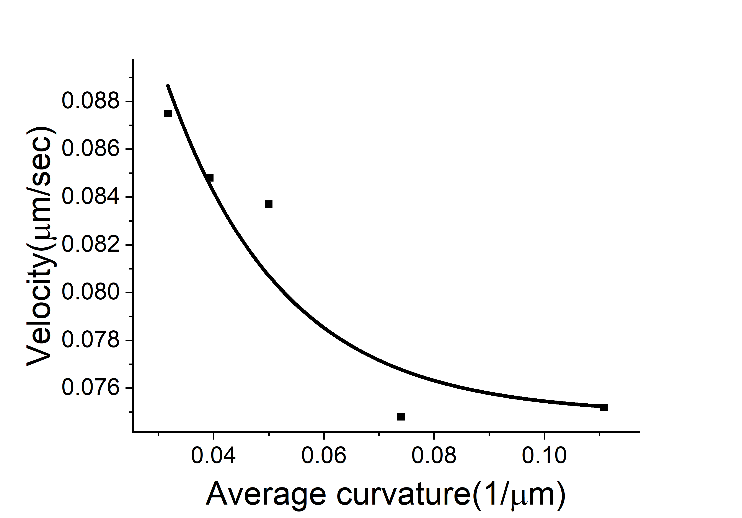

**I**

**II**

**A	1	2**

**B**

**C**

**Supplementary Fig 5**. (A) 1.) Relation between the wavefront velocity and average curvature from the experimental data for the centrifugal wavefront. A 2.) Nonlinear relationship between the wavefront velocity and average curvature for the centripetal wavefront. (B) Exponential fit for stage I of the centripetal wavefront travel as shown in (A 2). (C) Exponential fit for stage II of the centripetal wavefront travel shown in (A 2).

**Mathematical model of membrane dynamics in single cell *E. coli.***

We formulated a conductance model to explain the membrane potential dynamics in single *E. coli* cells. Our biophysical model is of the type originally used by Hodgkin and Huxley to explain excitability in squid axons [1]. We extended a version of this model which was previously used to study metabolic stress in *B. subtilis* biofilm [2,3],

$\frac{dV}{dt}=-g_{Q}z^{4}\left( V-V_{Q} \right)-g_{Kch}n^{4}\left( V-V_{Kch} \right)-g_{L}\left( V-V_{L} \right), 1$

where $g_{Q}$, $g_{Kch}$ and $g_{L}$ represent the conductance of the $Q^{+}$, $K^{+}$ and the leak channel respectively. $V_{q}$ , $V_{Kch}$ and $V_{L}$ represents the Nernst potentials for the $Q^{+}$, $K^{+}$ and the leak ions respectively.

We assumed that both of the positively charged ion channels (Q and Kch) have four subunits which are their four activation gates. The variable $z$ represents the gating variable for the $Q^{+}$channels. The fraction of time the $Q^{+}$ channel is open can then be represented by

$\frac{dz}{dt}=\alpha\left( S \right)\left( 1-z \right)-\beta_{z}. 2$

The $Q^{+}$channel opening rate $\alpha\left( S \right)$ depends on the accumulation of the light induced stress. This stress is related to the levels of ROS. $\beta_{z}$ represents the rate at which open channels close.

The ion channel gating variable *n* incorporates the fraction of time the potassium channel $Kch$ is open. It has the relation

$\frac{dn}{dt}=\alpha\left( S \right)\left( 1-n \right)-\beta_{n}, 3$

where $\alpha\left( S \right)$ stands for the opening rate for the $K^{+}$ and depends on the light-induced ROS stress. Blee *et al*  [3], has already shown that blue-light raises the level of ROS and causes the ion-channels to open.

The stress dependency of $\alpha\left( S \right)$ can be further described as

$$\alpha\left( S \right)=\frac{\alpha_{0}S^{r}}{S_{th}^{r}+S^{r}} , 4$$

where $S_{th}$is the threshold stress value for ion channel opening, $r$ is the cooperativity factor and $\alpha_{0}$ is the maximal opening rate.

Motivated by the description of neuronal excitability [1] and the works on bacterial signaling dynamics [2,3], we assumed that the ROS-induced stress induced by blue-light irradiation is related to the action potential in *E. coli* cells,

$\frac{ds}{dt}=\frac{\propto_{s}(V_{th}-V)}{exp\left( \frac{V_{th}-V}{\sigma}-1 \right)}-\gamma_{s}S . 5$

The term $\frac{\propto_{s}(V_{th}-V)}{exp\left( \frac{V_{th}-V}{\sigma}-1 \right)}$ is a threshold-linear function of the *E. coli* cell membrane hyperpolarisation, $\alpha_{s}$ is the stress production constant and $\gamma_{s}$ is the stress decay rate. We assume that the functions $\alpha_{s}$ and $\gamma_{s}$ have a strong link to the gating of the ion channels.

Following Prindle et al  [2], we assumed that the reversal potential increases linearly with the excess extracellular positively charged ion concentration and the relationship between the extracellular ion concentration and the reversal potential can be stated as

$\frac{dE}{dt}=F_{g_{Q}}z^{4}\left( V-V_{Q} \right)+F_{g_{k}}n^{4}\left( V-V_{K} \right)-\gamma_{e}E , 6$

where $E$ represent the extracellular ion concentration. The first and second term on the right-hand-side expresses the link between the extracellular $q^{+}$ and $K^{+}$ions, and the membrane potential. $F_{g_{Q}}$ and $F_{g_{k}}$ incorporates capacitance of the membrane to each of the ion channels.

Following Prindle, we represent the inverse relationship between the ThT concentration and the efflux of the positively charged ions (hyperpolarization) by a linear function,

$\frac{dH}{dt}=\alpha_{t}\left( V_{0}-V \right)-\gamma_{t}H$	. 7

The ThT decay rate is represented by $\gamma_{t}$ and $\alpha_{t}$ is the ThT uptake rate coefficient. $V$ is the membrane potential at a instant of time while $V_{0}$ is the membrane potential in the absence of the ROS (the resting potential).

Agent-based Fire-Diffuse-Fire Model

The fire-diffuse fire (FDF) model was previously developed to study intracellular calcium dynamics in living cells [4,5]. In the model, once an individual cell at a particular site in the biofilm reaches a threshold concentration $c^{*}$, the cells fires, releasing a given concentration $\sigma$ of the $C^{+}$ions. $C^{+}$incorporates both unknown ions and potassium ions ${(K}^{+})$ involved in our spike initiation and propagation in *E. coli* biofilms. A $C^{+}$ wavefront propagates within cells in the biofilm due to progressive firing of neighboring cells due to $C^{+}$ released close by.

The FDF model is based on a type of reaction-diffusion equation and can be written in 3 dimensions as

$$\frac{\partial C}{\partial t}=D_{C}\nabla^{2}C+ \sigma\sum_{i} \delta\left( \hat{r}-r_{i} \right)\delta\left( t-t_{i} \right) , 8$$

where $C$ is the chemical field concentration, $D_{C}$ is the diffusion coefficient of the ions, $\sigma$ is the fixed amount of chemical $C$, $b$ is the spacing between release sites, $t_{i}$ is the time the chemical attains threshold value $c^{*}$ at the $i^{th}$ release site and $\delta$ is the Dirac delta.

This original formulation of the FDF model does not include a decay term and results in the dynamics of the ion rising monotonically  [6]. To correctly reflect findings in our experiment, we include a decay term, $-\vartheta C$, which allows for refractoriness of the system.

$$\frac{\partial C}{\partial t}=D_{C}\nabla^{2}C-\vartheta C+ \sigma\sum_{i} \delta\left( \hat{r}-r_{i} \right)\delta\left( t-t_{i} \right) . 9$$

The first term $D_{C}\nabla^{2}C$ on the right-hand-term represent the diffusion term, while the remaining term on the right-hand-side, $-\vartheta C+ \sigma\sum_{i} \partial\left( \hat{r}-r_{i} \right)\partial\left( t-t_{i} \right)$ are the reaction terms. $\vartheta$ is the ion decay rate.

The solution of equation 9 in one dimension for bacteria placed on a perfect lattice $b$ is

$$C_{i}\left( x,t \right)=\sigma\frac{H(t-t_{i})}{\sqrt{4\pi D_{C}(t-t_{i})}}\exp\left( -\frac{\left( x-ib \right)^{2}}{4D\left( t-t_{i} \right)}-\vartheta\left( t-t_{i} \right) \right) , 10$$

where $H(t-t_{i})$ is the Heaviside function so the cells at the $i^{th}$ position only fires at the time $t_{i}$. $C_{i}\left( x,t \right)$ is the ion concentration for the individual cell (cell $i$).

For all cells along the x-axis, we have

$$C\left( x,t \right)=\sum_{i} \sigma\frac{H(t-t_{i})}{\sqrt{4\pi D_{C}(t-t_{i})}}\exp\left( -\frac{\left( x-ib \right)^{2}}{4D\left( t-t_{i} \right)}-\vartheta\left( t-t_{i} \right) \right) , 11$$

where

$C\left( x,t \right)=\sum_{i} C_{i}\left( x,t \right) . 12$

Suppose we assume that cells at definite location, say $i=N, N-1,$ have spiked at definite times $t_{N}>t_{N}-1>\ldots$, it is expected that they could only spike again at a time $t_{N+1}$ . This time $t_{N+1}$ is determined by when the field concentration $C$, at time $t_{N+1}$ reaches the threshold value $q^{*}$. Therefore, we expect a steady wave to propagate only when the neighboring cells exhibit a constant time difference, $\tau=t_{i}- t_{i-1}$ , between their firing. This time constant becomes a solution of the equation

$$\frac{c^{*}b}{\sigma}=\sum_{n=1}^{\infty} \frac{1}{\sqrt{4\pi n\rho}}exp\left( \frac{-n}{4\rho}-\eta^{2}n \right), 13$$

where $\rho=\frac{D_{C}\tau}{b^{2}}$ and $\eta^{2}=\frac{\vartheta b^{2}}{D_{C}}$ are both dimensionless. A wave propagates when $\frac{c^{*}b}{\sigma}<1$. The velocity $\nu$of the propagating wave can then be obtained from $\frac{b}{\tau}$ .

While an exact solution of this 1D form of the FDF model with regularly placed bacteria can be obtained as shown, we used agent-based modelling to solve this model in three-dimensions with heterogeneously placed bacteria i.e. to solve equation 9. We had used the FDF model to successfully model electrical signalling in 2D growing biofilm [7]. We adopted and applied the conditions and rules of this FDF model in our agent-based modelling of three-dimensional biofilm. A key extension was incorporating it in the agent-based modelling tool, BSim, to simulate electrical signalling in three-dimensional biofilms.

BSim is an open-source customizable Java based agent-based modelling software. It is specifically built to incorporate the effects of complex spatial environmental features and community heterogeneity [8]. We modelled the microfluidic device as a three-dimensional fluid in a rectangular environment. We implemented the biologically relevant parameters and field constants (See Table 1) to obtain similar ion-channel mediated waves to those observed in three-dimensional biofilms. The positively charged ions $Q^{+},$ were described via their diffusion coefficients.

A BSim environment is built as an octree which also controls the spread of the chemical in the $x,y, z$space. A Bsim environment, $64\times64\times64$ microns in size with fixed boundaries was created (Fig 7A). A spherical biofilm of radius $32 \mu m$ containing $\sim6000$cells was implemented. The signalling ions chemical has a diffusivity and decay rate of $0.02 {\mu m}^{2}/s$ and $7\times{10}^{-3} {molecules}^{-1}$ respectively.

The ion diffusion inside the BSim space was calculated according to Fick’s law of diffusion,

$\frac{\partial C}{\partial t}=D\nabla^{2}C, 14$

where $C$ is the ion concentration in the field and $D$ is the diffusion coefficient of the ions. Java constructors were used to initialize the fields of ion concentration and control their diffusivity and decay rate.

**Table S1**

| **Parameter** | **Description** | **Value** | **Units** |
| --- | --- | --- | --- |
| $g_{K}$ | Potassium channel conductance | $90$ | ${min}^{-1}$ |
| $g_{L}$ | Leak channel conductance | $0.2$ | ${min}^{-1}$ |
| $g_{q}$ | $q$ channel conductance | $5$ | ${min}^{-1}$ |
| $V_{K}$ | Nernst Potential for Potassium | $-94$ | $mV$ |
| $V_{q}$ | Nernst potential for $q$ channel | $-200$ | $mV$ |
| $V_{L}$ | Nernst potential for leak channel | $-156$ | $mV$ |
| $S_{th}$ | Stress threshold for opening of ion channels | $0.04$ | $\mu M$ |
| $V_{th}$ | Voltage threshold for stress production | $-150$ | $mV$ |
| $a0$ | Maximum rate of channel opening | $2$ | ${min}^{-1}$ |
| $b0$ | Channel opening rate decay constant | $1.3$ | ${min}^{-1}$ |
| $r$ | Cooperativity parameter for ion channels | $1$ | $-$ |
| $\sigma$ | Stress threshold sharpness coefficient | $0.2$ | $mV$ |
| $y_{e}$ | Extracellular ion relaxation rate | $10$ | ${min}^{-1}$ |
| $dl$ | Leak slope coefficient | $8$ | $mV/mM$ |
| $d_{q}$ | $q$ channel slope coefficient | $2$ | $mV/mM$ |
| $d_{K}$ | Potassium channel slope coefficient | $1$ | $mV/mM$ |
| $\alpha_{s}$ | Stress production slope coefficient | $0.001$ | $\mu M/(\min mV)$ |
| $\alpha_{t}$ | ThT uptake rate coefficient | $0.4$ | $\mu M/(\min mV)$ |
| $\gamma_{s}$ | Stress decay rate | $0.1$ | ${min}^{-1}$ |
| $\gamma_{t}$ | ThT decay rate | $4$ | ${min}^{-1}$ |
| $F$ | Membrane capacitance | $5.6$ | $mM/mV$ |

**Table S1: Parameters for the Mathematical modeling of membrane potential dynamics in *E. coli.***

**Table S2**

| **Field and model parameters** | **Symbols** | **Values** |
| --- | --- | --- |
| Diffusion coefficient | $D_{C}$ | $0.1 {\mu m}^{2}/s$ |
| Simulation time step | $T_{s}$ | $1 sec$ |
| Total time of simulation for the 2 peaks | $t_{s}$ | $3660 sec$ |
| Total time of simulation for the first peak | $t_{s}$ | $720 sec$ |
| Size of fluid-filled environment | $S_{b}$ | $64 \mu m\times64 \mu m\times64 \mu m$ |
| Fixed amount of chemical added | $\sigma$ | $5\times{10}^{9}{\mu m}^{-3}$ |
| Threshold concentration to fire | $c^{*}$ | ${10}^{3}{\mu m}^{-3}$ |
| Decay rate (for refractoriness) | $\vartheta$ | $7\times{10}^{-3} {molecules}^{-1}$ |

**Table S2: Parameters used for the Agent-based Fire-diffuse-fire Model in the three-dimensional *E. coli* biofilm.**

**Table S3**

| **Fit constants for the centrifugal wavefront** | **Values** |
| --- | --- |
| $\boldsymbol{V}_{\boldsymbol{0}}$ | $0.23\pm0.04$ |
| $\boldsymbol{A}$ | $0.010\pm0.008$ |
| $\boldsymbol{z}$ | $0.020\pm0.002$ |
| **Fit constants for the centripetal wavefront – Stage I** |  |
| $\boldsymbol{V}_{\boldsymbol{0}}$ | $0.074\pm0.001$ |
| $\boldsymbol{A}$ | $-$ |
| $\boldsymbol{z}$ | $0.018\pm0.002$ |
| **Fit constants for the centripetal wavefront – Stage II** |  |
| $\boldsymbol{V}_{\boldsymbol{0}}$ | $0.075\pm0.003$ |
| $\boldsymbol{A}$ | $-$ |
| $\boldsymbol{z}$ | $-0.02\pm0.01$ |

**Table S3: Fit constants for the centrifugal and centripetal wavefronts from experimental data**

**Table S4:** Media recipe, bacterial strains, software and microfluidics components.

| **Media** | **Recipe** |  |
| --- | --- | --- |
| Luria Broth (LB) | 10g/l NaCl, 5g/l yeast extract, 10g/l Tryptone. Distilled water to 1L |  |
| LB agar | 10g/l NaCl, 5g/l yeast extract, 10g/l Tryptone, 15 g/l agar. |  |
| Minimal Media (M9) (1L) | M9 salts (64g/l Na_2_HPO_4_·7H_2_0, 15g/l KH_2_PO_4_, 2.5g/l NaCl, 5.0g NH_4_Cl), 1M CaCl_2_, 1M MgSO_4_, 10% Arginine, 20% of glucose |  |
| **Experimental Models: Organisms/Strains** | | |
| *E. coli* DH5α | Ian Robert’s lab |  |
| *E. coli* DH5α (∆*Kch*) | This study |  |
| *E. coli* DH5α (∆MscK) | This study |  |
| *E. coli* DH5α (∆MscL) | This study |  |
| *E. coli* DH5α (∆MscS) | This study |  |
| *E. coli* BW25113 (JW1242-1) | Keio Collection  [9] |  |
| *E. coli* BW25113 (∆*Kch*) | Keio Collection  [9] |  |
| *E. coli* BW25113 (∆MscL) | Keio Collection  [9] |  |
| *E. coli* BW25113 (∆MscS) | Keio Collection  [9] |  |
| *E. coli* BW25113 (∆MscK) | Keio Collection  [9] |  |
| **REAGENT or RESOURCE** | **SOURCE** | **IDENTIFIER** |
| Propidium Iodide (PI) | ThermoFisher Scientific | Cat #: P1304MP |
| Thioflavin T (ThT) | Sigma-Aldrich | Cat #: T3516 |
| Carbonyl cyanide 3-chlorophenylhydrazone (CCCP) | Sigma-Aldrich | Cat #: C2759 |
| ION Potassium Green–4 (IPG-4) AM | ION Biosciences | https://ionbiosciences.com/product/ipg-4-am/ |
| Tetramethylrhodamine methyl ester perchlorate (TMRM | Sigma-Aldrich | Cat No: T5428 |
| pkLL11-GCaMP6f bb100 | A gift from Joel Kralj | Addgene plasmid #158983 |
| **Software and Algorithms** | | |
| ImageJ Fiji | [10] | https://fiji.sc/ |
| Python (Anaconda) | Python | https://anaconda.org/ContinuumIO |
| MATLAB | MATLAB | <https://uk.mathworks.com>  /products/matlab.html |
| BiofilmQ | [11] | <https://drescherlab.org/data/biofilmQ/>  docs/usage/installation.html |
| OriginPro | https://www.originlab.com/origin | https://www.originlab.com/origin |
| IMARIS | https://imaris.oxinst.com/ | https://imaris.oxinst.com/ |
| **Microfluidics** | | |
| IBIDI flow cell | Thistle Scientific | Cat #: 80607 |
| IBIDI Elbow Luer Connector Male | Thistle Scientific | Cat #: 10802 |
| C-Flex laboratory tubing with I.D. x O.D. 1/32 in. x 3/32 in | Sigma-Aldrich | Cat #: T8413-25FT. |

**Table S4 continues:** Media recipe, bacterial strains, software and microfluidic components.
